# Multiomics data analysis using tensor decomposition based unsupervised feature extraction --Comparison with DIABLO--

**DOI:** 10.1101/591867

**Authors:** Y-h. Taguchi

## Abstract

Multiomics data analysis is the central issue of genomics science. In spite of that, there are not well defined methods that can integrate multomics data sets, which are formatted as matrices with different sizes. In this paper, I propose the usage of tensor decomposition based unsupervised feature extraction as a data mining tool for multiomics data set. It can successfully integrate miRNA expression, mRNA expression and proteome, which were used as a demonstration example of DIABLO that is the recently proposed advanced method for the integrated analysis of multiomics data set.

## 1 Introduction

Multiomics data, including miRNA expression, mRNA expression, promoter methylation and histone modification, have recently come to be measured over the various biological problems. In contrast to the rapid development of measurement technology, the data analysis pipeline developed very slowly. This is principally because we have never faced the flood of data set until very recently; measuring data set has ever been more expensive than analyzing it. Thus, only limited efforts have been spent for analysis of high dimensional data that has ever been rarely available.

Multiomics data is a typical high dimensional dataset; the number of samples (typically <10^2^) is always less than number of features, i.e., that of mRNAs (~10^4^), that of miRNAs (~10^3^) and that of methylation sites (~10^5^). There are several methods proposed in order to integrate multiomics data formatted as matrices with distinct sizes. **Fig. 1** shows typical three strategies to integrate three matrices that share the *M* samples for which three distinct features whose numbers are *N*_1_, *N*_2_, and *N*_3_ are measured. **Fig. 1**(A), contraction, is simply merging three matrices such that they share *M* samples as rows while *N*_1_, *N*_2_, and *N*_3_ features are aligned as columns. After merging them, all downstream analyses are performed with assuming only one feature whose number is *N*_1_+*N*_2_+*N*_3_. Although this strategy looks simple, there can be multiple disadvantages. First of all, if the number of individual features, *N*_1_, *N*_2_, and *N*_3_, differ from one another so much, features having the smallest number might be neglected.

**Fig. 1.**
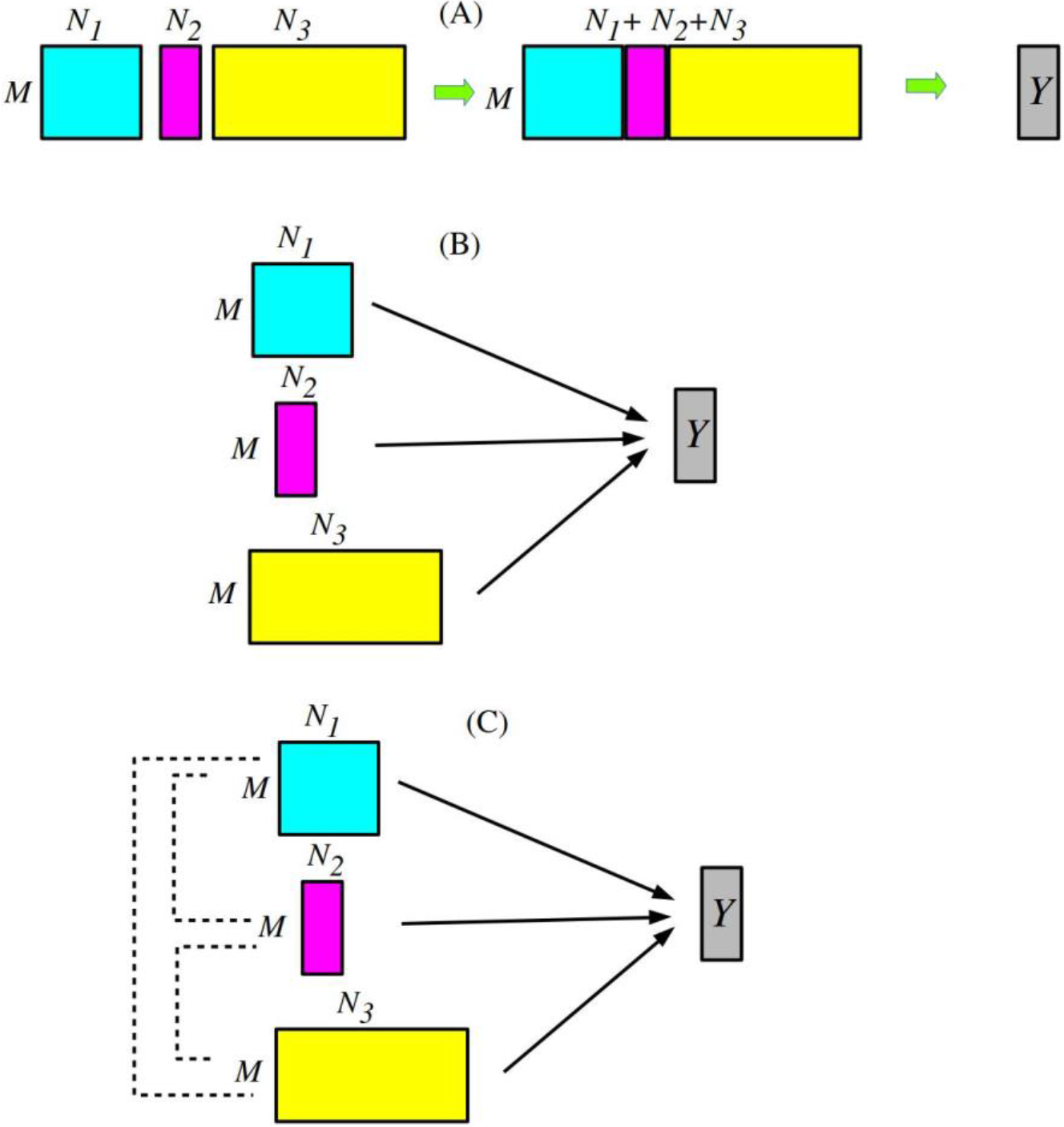
Three distinct strategies that integrate multiomics formatted as matrices with distinct dimensions, *M* × *N*_1_, *M* × *N*_2_, and *M* × *N*_3_. (A) Contraction (B) Ensemble (C) DIABLO.

This might prevent us from considering three features equally. Even if the numbers are almost equal, if some feature has much larger amplitude than others, it also prevents us from dealing with them equally. In this sense, it is very usual to normalize individual features before the contraction. Nevertheless, the normalization might also affect the outcome, because how to normalize them is quite subjective.

In contrast to contraction, ensemble strategy shown in **Fig. 1**(B) oppositely does not integrate multiomics data until very last stage. The simplest ensemble strategy is to train individual feature independently, and decision is made upon the majority rule. This strategy is also simple because we do not need any new strategies considering integration of multi omics data set. The disadvantage of this strategy is obvious. There is no justification for giving one vote to individual feature. If the majority of features are useless, i.e., do not have any practical relationship with outcome *Y*, voting system simple increases the noise. In this sense, ensemble can be worse than contraction that can disregard useless features during training.

The third strategy shown in **Fig. 1**(C), DIABLO [1], which is a part of mixOmics [2] that aims integrated analysis of multiomics data, is more advanced method. In DIABLO strategy, not individual features but sets (pairs) of individual features are employed for training. Then, outcome *Y* is predicted based upon ensemble strategy of them. Thus, DIABLO strategy has more ability to learn effective features from available multiomics data. One disadvantage is that how to relate individual features must be designed by human beings; this process can be quite subjective.

In this paper, in contrast to these three strategies in **Fig. 1**, unsupervised strategy based upon tensor decomposition (TD), in short, “TD based unsupervised feature extraction (FE)”, is proposed and is applied to multiomics data set. As can be seen later, TD based unsupervised FE achieves performance competitive with that achieved by DIABLO strategy.

## 2 Materials and Methods

### 2.1 Multiomics data set

Data set used here is included in mixOmics package^1^. As described in the web page “Case study: TCGA”^2^, this data set can be loaded into R using data(‘breast.TCGA’) command after installing mixOmics package. It includes 150 samples composed of 45 Basal, 30 Her2 and 75 LumA subtypes, to which 200 mRNAs, 184 miRNAs and 142 proteome expression are measured.

### 2.2 Tensor decomposition

We apply TD with higher order singular value decomposition (HOSVD) [3] algorithm to case I type I tensor [4]. Starting from three matrices, 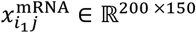, 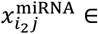 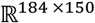, and 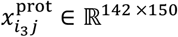, we generate four mode tensor

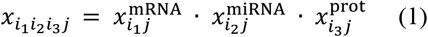

and HOSVD is applied it as

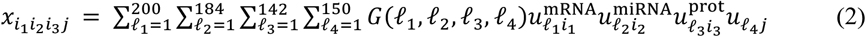

where 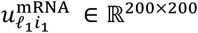, 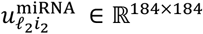, 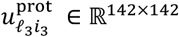, and 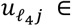 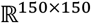, are singular value matrices (they are also orthogonal matrices) and G ∈ ℝ^200 ×184 ×142 ×150^ is a core tensor.

## 3 Results

### 3.1 DIABLO

Here I briefly summarize how well DIABLO works when it is applied to data set described in Sec. 2.1 as denoted in the web page “Case study: TCGA”^2^. The interaction assumed is mRNA-miRNA, mRNA-proteome, and miRNA-proteome.

1. Three subtypes can be discriminated with the success rate of 95 % with using more than two components generated by DIABLO.
2. 15 mRNAs, 18 miRNAs and 7 proteins are selected.
3. Using features selected in step 2, heatmap is drawn. These three subtypes are well separated in the heatmap. Thus, DIABLO successfully selects limited number of features that discriminate three subtypes.

Thus, the point is if TD based unsupervised FE can achieve similar performance as those by DIABLO or not.

### 3.2 TD based unsupervised FE

As described in Sec. 2.2, HOSVD is applied to case I type I tensor, eq. (1). Then two singular value vectors, *u*_1*j*_ and *u*_4*j*_, turn out to discriminate three subtypes well (**Fig. 2**). In order to quantitatively evaluate how well these two singular value vector discriminates three subtypes, we apply linear discriminant analysis (LDA) with using leave one out cross validation (LOOCV) (**Table 1**). It achieved as high as 95 % accuracy, which is almost equal to that by DIABLO.

**Fig. 2.**
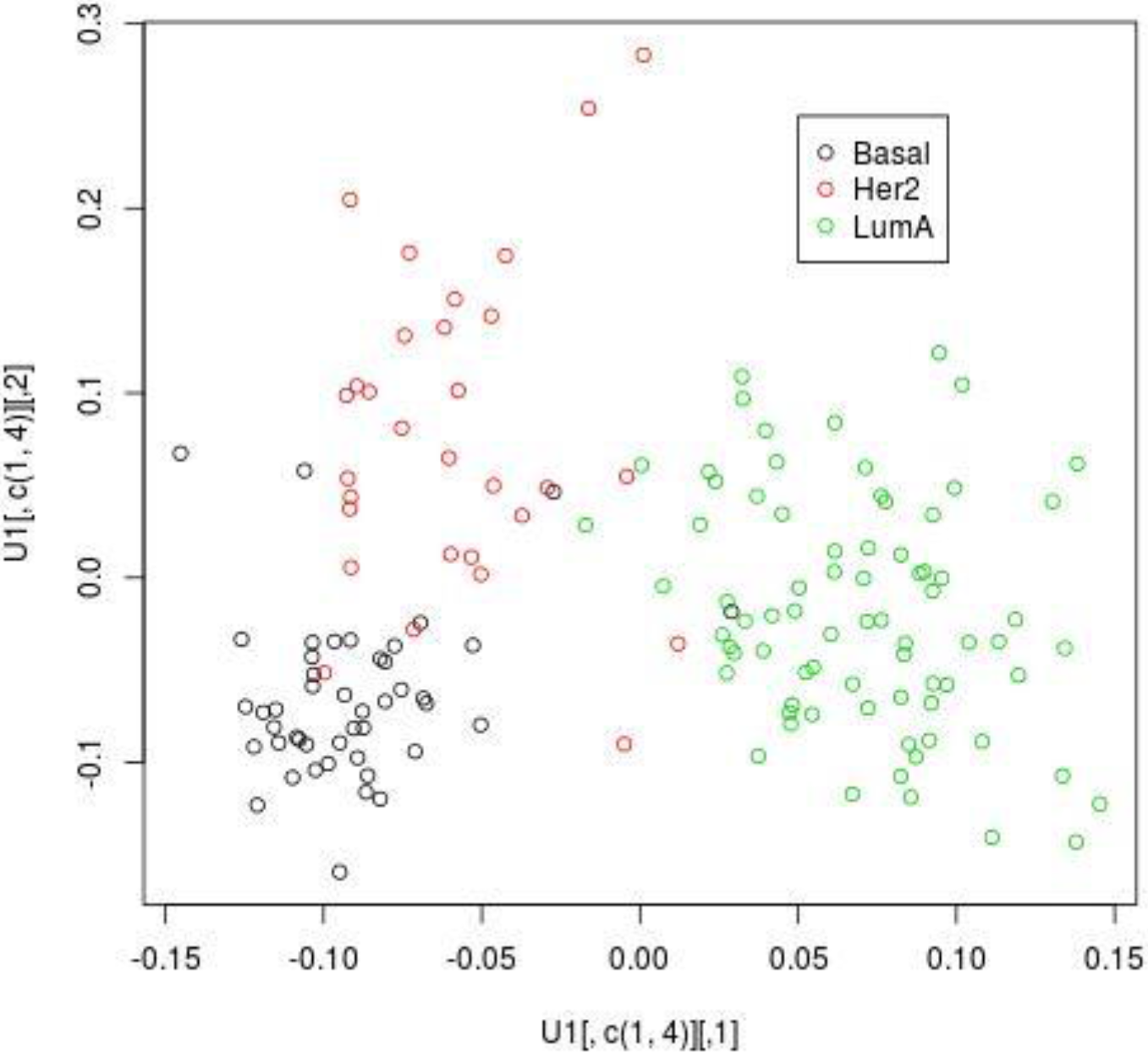
Scatter plot of *u*_1*j*_ (horizontal axis) and *u*_4*j*_ (vertical axis).

**Table 1.** The confusion table obtained by applying LDA to three cancer subtypes with *u*_1*j*_ and *u*_4*j*_. LOOCV is used. Row: inference, column: true subtypes. Numbers in bold represent those of correctly predicted samples.

|  | Basel | Her2 | LumA |
| --- | --- | --- | --- |
| Basel | **42** | 4 | 0 |
| Her2 | 2 | **25** | 2 |
| LumA | 1 | 1 | **73** |

Next, in order to see if the limited number of mRNAs, miRNAs and proteomes can be selected to discriminate three cancer subtypes well, we select subsets of these three features. In order that, we first need to find which 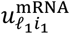, 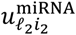, and 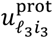 are most associated with *u*_1*j*_ and *u*_4*j*_. This requires the identification of core tensor, *G*(*ℓ*_1_, *ℓ*_2_, *ℓ*_3_, *ℓ*_4_), having the largest absolute values with *ℓ*_4_ = 1,4. **Table 2** shows the list of those *G*(*ℓ*_1_, *ℓ*_2_, *ℓ*_3_, *ℓ*_4_) s with the descending order of absolute values of *G*(*ℓ*_1_, *ℓ*_2_, *ℓ*_3_, *ℓ*_4_)s. It is obvious that *G*(*ℓ*_1_, *ℓ*_2_, *ℓ*_3_, *ℓ*_4_)s with 1 ≤ *ℓ*_1_, *ℓ*_2_ ≤ 2 and 1 ≤ *ℓ*_3_ ≤ 4 have larger absolute values. Then we compute the squared summation of singular value vectors attributed to *i*_1_th mRNA, *i*_2_th miRNA, and *i*_3_th proteome as

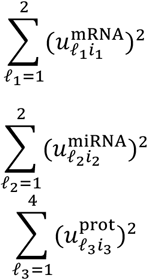

and select top ranked ten mRNAs, miRNAs and proteome with larger squared summation of singular value vectors.

**Table 2.** *G*(*ℓ*_1_, *ℓ*_2_, *ℓ*_3_, *ℓ*_4_)s sorted descending order of absolute values among those having *ℓ*_4_ = 1, 4.

| rank | $\ell_1$ | $\ell_2$ | $\ell_3$ | $\ell_4$ | $G(\ell_1, \ell_2, \ell_3, \ell_4)$ |
| --- | --- | --- | --- | --- | --- |
| 1 | 1 | 1 | 1 | 1 | -407857.582 |
| 2 | 1 | 1 | 4 | 4 | -209720.615 |
| 3 | 2 | 1 | 1 | 4 | -20452.480 |
| 4 | 2 | 1 | 3 | 1 | -11677.505 |
| 5 | 2 | 1 | 4 | 1 | -10428.742 |
| 6 | 2 | 1 | 2 | 1 | 10157.467 |
| 7 | 1 | 1 | 2 | 1 | -8973.774 |
| 8 | 1 | 2 | 1 | 4 | 8360.976 |
| 9 | 2 | 1 | 5 | 4 | -6628.467 |
| 10 | 1 | 1 | 3 | 4 | 6623.046 |

**Fig. 3**. shows the heatmap generated using selected 10 mRNAs, 10 miRNAs and 10 proteome. It is obvious that three subtypes are clustered well separately; it is competitively well as compared with the heatmap generated by DIABLO^2^. In addition to this, three features that share same profiles are clustered together as in the result by DIABLO (see hierarchical clustering of columns in **Fig. 3**). This suggests that TD based unsupervised FE has ability to select distinct features sharing same profiles over three subtypes as DIABLO achieved.

**Fig. 3.**
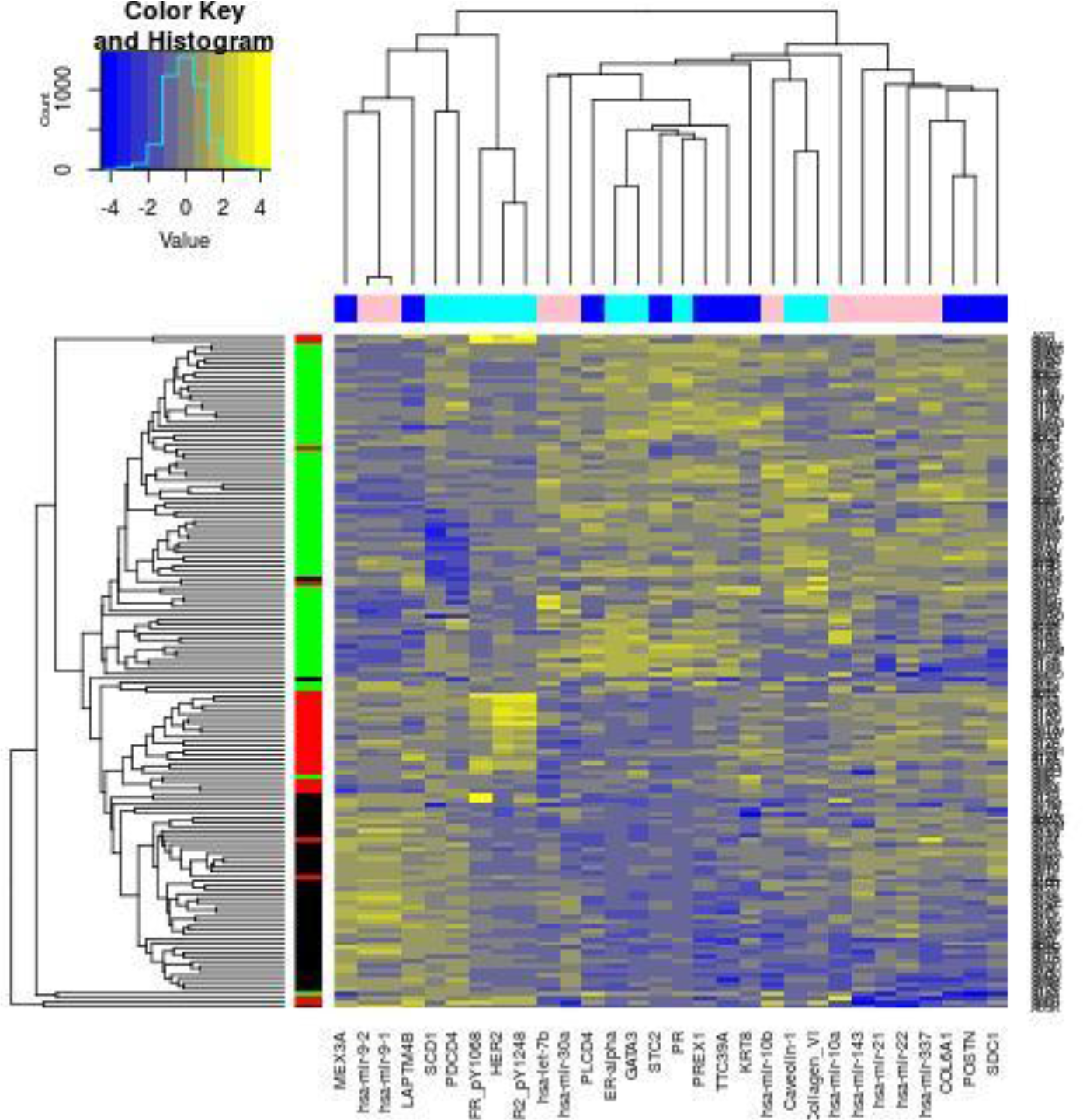
Heatmap of selected 10 mRNAs (blue), 10 miRNAs (pink), and 10 proteome (cyan) .in columns and 150 samples (black:Basel, red:Her2, green:Luma). Yellow: expressed expression, blue: depressed expression.

## 4 Discussions

In this paper, I applied recently proposed TD based unsupervised FE, which was applied to wide range of studies [4–35], to multiomics data analysis. Apparently, the performance achieved by TD based unsupervised FE is at most competitive to that of DIABLO. Thus, one might wonder why TD based unsupervised FE is recommended in this study. There are multiple reasons for this recommendation. First, DIABLO used random number seed to perform analysis. This means, every time we select different random number seed, we inevitably have distinct sets of selected mRNAs, miRNAs and proteome, although overall performance remains unchanged. This might be problematic because we might be interested in identification of disease causing genes. If distinct set of features are selected every time we change random number seed, it might prevent us from interpreting the outcome biologically. In contrast to this, TD based unsupervised FE can give us quite stable outcomes. Not only TD based unsupervised FE does not need random number seed, but also it is quite stable even when samples are resampled [4–35]. In this point, TD based unsupervised FE is more suitable to be employed than DIABLO form the biological point of views.

Also from the computational point of views, TD based unsupervised FE is recommended more than DIABLO. From the point of computational time, DIABLO requires more time than TD based unsupervised FE, because DIABLO needs to learn from the data set and labeling while TD based unsupervised FE does not require this process due to unsupervised nature. In this sense, if these two achieve equally, TD based unsupervised FE is more recommended method than DIABLO.

## Appendix R code for TD based unsupervised FE

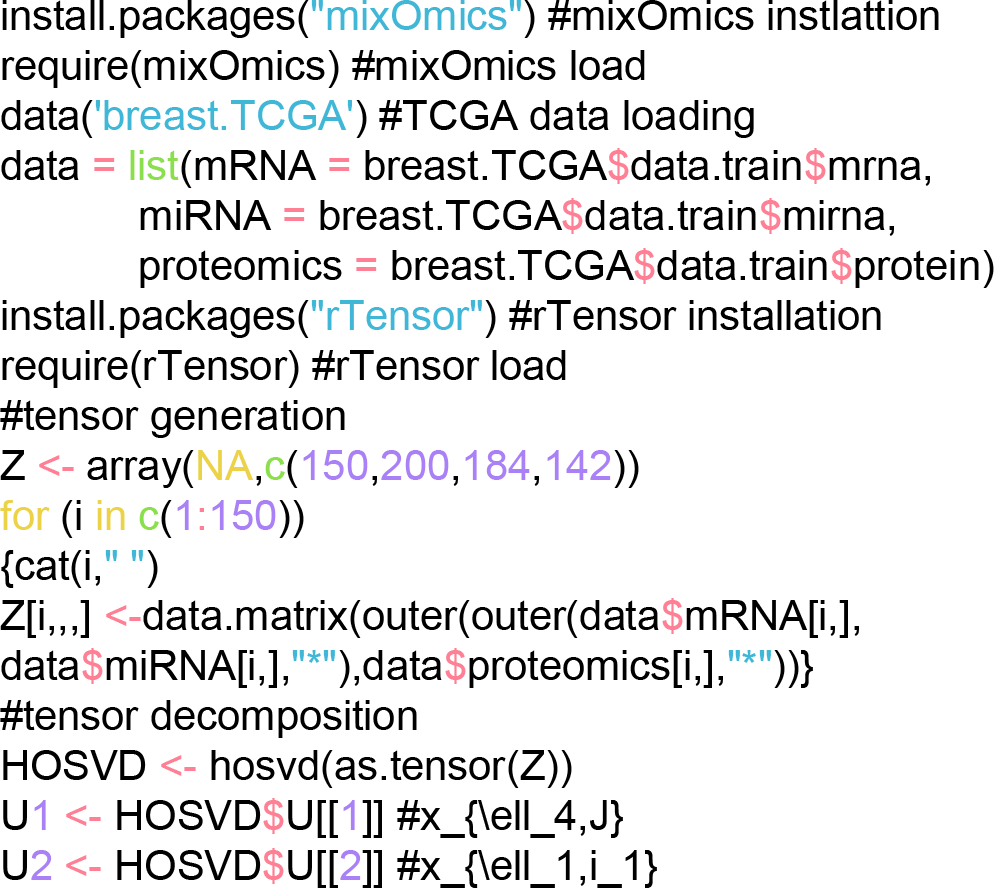

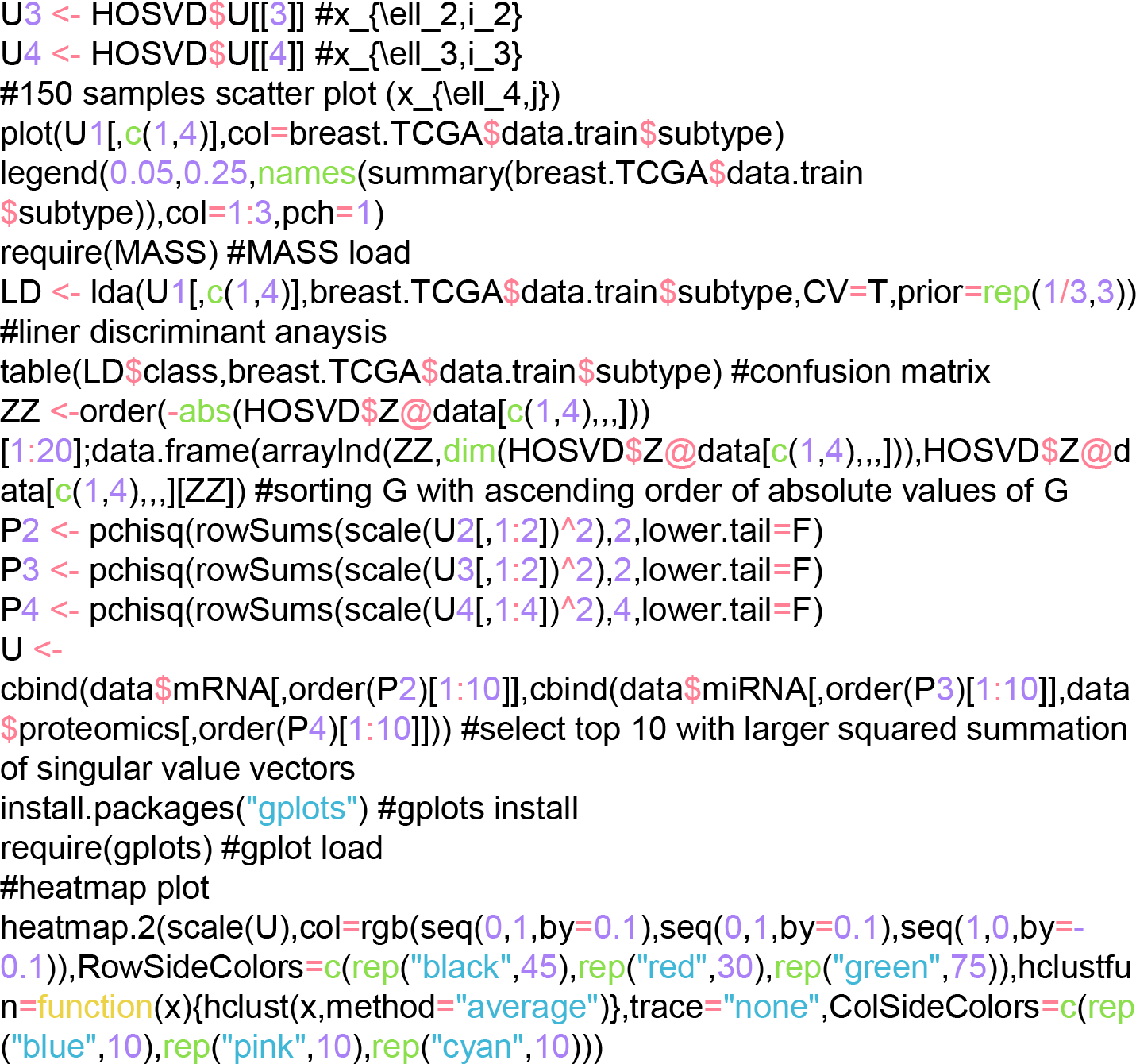

## Footnotes

1 http://www.bioconductor.org/packages/release/bioc/html/mixOmics.html

2 http://mixomics.org/mixdiablo/case-study-tcga/

